# Defining marine bacterioplankton community assembly rules by contrasting the importance of environmental determinants and biotic interactions

**DOI:** 10.1101/2022.05.02.490371

**Authors:** Michael P. Doane, Martin Ostrowski, Mark Brown, Anna Bramucci, Levente Bodrossy, Jodie van de Kamp, Andrew Bissett, Peter Steinberg, Martina A. Doblin, Justin Seymour

**Author notes:** Michael P. Doane –. Martin Ostrowski –. Mark Brown –. Anna Bramucci –. Levente Bodrossy –. Jodie van de Kamp. Andrew Bissett –. Martina A. Doblin –. Justin Seymour –.

## Abstract

Bacterioplankton communities play major roles in governing marine productivity and biogeochemical cycling, yet what drives the relative influence of the types of deterministic ecological processes which result in diversity patterns remains unclear. Here we examine how differing deterministic processes (environmental factors and biotic interactions) drive temporal dynamics of bacterioplankton diversity at three different oceanographic time-series locations, spanning 15 degrees of latitude, which are each characterized by different environmental conditions and varying degrees of seasonality. Monthly surface samples, collected over a period of 5.5 years, were analyzed using 16S rRNA amplicon sequencing. The high and mid-latitude sites of Maria Island and Port Hacking were characterized by high and intermediate levels of environmental heterogeneity respectively, with both alpha (local) diversity (72 % and 24 % of total variation) and beta diversity (32 % and 30 %) patterns within bacterioplankton assemblages primarily explained by environmental determinants, including day length, ammonium, and mixed layer depth. In contrast, at North Stradbroke Island, a sub-tropical location where environmental conditions are less seasonally variable, interspecific interactions were of increased importance in structuring bacterioplankton diversity (alpha diversity: 33 %; beta diversity: 26 %) with environment only contributing 11 and 13 % to predicting diversity, respectively. Our results demonstrate that bacterioplankton diversity is the result of both deterministic environmental and biotic processes and that the importance of these different deterministic processes varies, potential in response to environmental heterogeneity.

**Importance:** Marine bacterioplankton drives important biological processes, including the cycling of key nutrients or fixing atmospheric carbon. Therefore, to predict future climate scenarios its critical to model these communities accurately. Processes that drive bacterioplankton diversity patterns in the oceans however remain unresolved, with most studies focusing on deterministic environmental drivers, ie temperature or available inorganic nutrients. Biotic deterministic processes including interactions among individuals are also important for structuring diversity patterns, however, this is rarely included to predict bacterioplankton communities. We develop an approach for determining the relative contribution of environmental and potential biotic interactions that structure marine bacterioplankton at three series at different latitudes. Environmental factors best predicted temporal trends in bacterioplankton diversity at the two high latitude time series, while biotic influence was most apparent at the low latitude time series. Our results suggest environmental heterogeneity is an important attribute driving the contribution of varying deterministic influence of bacterioplankton diversity.

## Introduction

Bacterial community structure influences ecosystem function in fundamental ways across all natural environments [1–3], including the ocean [4], where microorganisms represent the base of the food web and are the principal mediators of biogeochemical cycles [5]. Ecological diversity underscores community structure, therefore, elucidating the processes that govern bacterioplankton diversity is critical for predicting marine ecosystem productivity and function. There are two alternative perspectives for how bacterial diversity assembles [6, 7]. One view is of determinism, where species are regulated by niche processes such as environmental filtering [8] stemming from physico-chemical factors such as inorganic nutrients availability and temperature [9, 10], as well as biotic interactions including competition, predator-prey or facultative interactions [11–13]. The other view is one of neutrality, where species are considered ecologically equivalent and therefore diversity consequently arise from stochastic birth, death, colonization, immigration and speciation [14–19]. The relative contribution of environmental, biotic interactions and stochastic processes, and how their importance changes over space and time is currently unresolved [20], meaning that the ability to interpret and predict marine bacterioplankton diversity is currently restricted.

In the ocean, both environmental factors and trophic interactions fundamentally govern bacterioplankton diversity [21, 22], in terms of both the number of co-occurring species (alpha diversity) and the commonality of species among environments or sampling points (beta diversity) [23, 24]. For instance, bacterioplankton community richness in the English channel, was highest during the winter months and strongly predicted by day length [25]. Landau et al., (2013) similarly found day length to strongly associate with marine bacterioplankton richness from temperate regions. In contrast, bacterioplankton community richness from the Antarctic region was negatively correlated with seasonal increases in chlorophyll-a (Chl-a), signaling potential interactions with algal blooms [26]. Community beta diversity patterns have also been shown to have environmental and biotic links. For instance, global samplings of surface bacterioplankton from the TARA dataset showed the strong effect of temperature and oxygen in driving community composition [27]. The San Pedro oceanographic time-series (SPOTS) in the California Bight, surface layer (0-5 m) bacterioplankton community beta diversity across 10 years was best predicted by abiotic factors including nitrate and day length change as well as Chl-a [28]. From a high-resolution coastal time-series, Needham et al. (2018) demonstrated with networks that bacterial abundance patterns were more strongly coupled to phytoplankton dynamics than other environmental factors, highlighting the role of biotic processes in structuring bacterioplankton community patterns. This study also noted the high correlated among groups of bacteria, indicating biotic interactions are not limited to vertical trophic interactions, but can occur horizontally through cross-feeding and antagonism, which are hypothesized to also fundamentally governing bacterioplankton diversity[22, 29, 30]. Similarly, bacteria-bacteria interactions were shown to be important for the maintenance of bacterioplankton diversity in the English Channel evidenced by bacterial OTUs having stronger correlation with other bacterial OTUs than with phytoplankton OTU’s and environmental factors [25]. Similarly, at SPOTS, network analysis demonstrated that bacteria, archaea, and eukaryotes had stronger correlation with one another than with any physico-chemical factors [31]. More recently, Lima-Mendez (2015), incorporated the abundance of eukaryotic and viral groups alongside environmental factors to demonstrate that abiotic factors explained a limited amount of direct variation in marine bacterioplankton diversity and that trophic and symbiotic interactions were significant contributors to overall diversity [32]. Collectively, these results underscore that while environmental factors are important regulators of bacterioplankton diversity, biotic interaction are apparent and potentially influence bacterioplankton more strongly at times, but the relative contribution of each deterministic type is yet to be resolved.

Marine environments are inherently dynamic in their environmental characteristics, fluctuating across scales of space (ie. micrometers to kilometers), and time (ie. microseconds to months) [21]. Therefore, distilling out specific factors responsible for diversity is particularly challenging, and may not accurately reflect the contemporary processes responsible for observed patterns [33]. Considering then the cumulative impacts of a set of environmental factors (or species interactions), and their relative contribution to diversity patterns is important because the impact magnitude of the ecological process is expected to vary in response to different ecological attributes (ie environmental heterogeneity) [20, 34, 35]. In one example, Langenheder et al. (2012) [36], showed that when environmental heterogeneity among rockpools was highest, beta diversity of bacterioplankton among the same rockpools was also highest, with deterministic processes emerging as the prevailing mechanism driving beta diversity; however when environmental heterogeneity among rockpools was low, and beta-diversity among rockpools was also relatively low, dispersal mechanisms became increasingly more important in driving beta diversity patterns.

Fluctuations in environmental heterogeneity have been demonstrated as important drivers of spatial beta diversity patterns in disparate ecosystems, including the Amazon river system [7] and soil bacteria communities [37, 38]. These results reveal that the relative contributions of deterministic processes in shaping spatial patterns bacterioplankton diversity can change through time in accordance with spatially distributed environmental heterogeneity [20, 39]; it remains to be shown however how deterministic influence is partitioned among environmental factors versus species interactions. In addition, marine bacterioplankton studies have focused primarily on understanding the influence of spatial environmental heterogeneity on bacterial diversity or temporal variability examined at a single location [20]. Therefore, current understanding of shifts in the relative importance of differing ecological processes on the temporal dynamics of bacterioplankton diversity at different locations is critically needed to understand if processes structing diversity patterns are universal across distinct environment or rather idiosyncratic to an environmental.

Distinguishing abiotic from biotic processes in structuring community diversity requires an effective means of identifying potential species interactions [40]. Herren and McMahon (2017)[41] developed a community complexity metric for phytoplankton microbial communities based on the product of the median correlation value of each organism in the dataset to its relative abundance value. The authors argue this value provided an index for quantifying the importance of potential interactions within the community. Indeed, their results demonstrated that characterizing the complexity of a community can improve the proportion of explained variation, but it remained unclear whether the explained variance was due to environmental and/or species interactions because the metric was based on correlations, which could arise due to multiple organisms independently tracking similar environmental factors. One potential means to overcome this limitation is by incorporating metrics that are applied to individual samples, derived from the number and strength of correlative interactions with other species in the community relative to environmental factors [32, 42]. A similar approach has been used to partition potential species interactions from environmental drivers in freshwater macro-organism communities, which highlighted the importance of species interactions in determining community structure [43].

Here, we use a 5.5 year time-series, including physico-chemical and 16s microbial community data to investigate the relative importance of environmental filtering versus inter-organismal interactions in influencing marine bacterioplankton structure. Three oceanographic time-series spanning 15° of latitude along the east Australian coastline allowed us to determine the relative contribution of environmental factors relative to potential biotic interactions. Bacterioplankton community structure was inferred by identifying the relative contribution of deterministic processes to shaping patterns of alpha and beta diversity. Our analysis involved the integration of a novel metric for inferring potential species interactions, defined as bacteria-bacteria and phytoplankton-bacterial interactions (biotic interactions), to discriminate among the relative importance of different deterministic processes in shaping bacterioplankton structure.

## Methods

### Reference station description and environmental data collection

Monthly surface water samples were collected from three oceanographic time-series stations located on the eastern continental shelf of Australia, as part of the Integrated Marine Observing System (IMOS) National Reference Station (NRS) monitoring program. These stations span latitudes of 27 to 42° S and include Maria Island (MAI: 42°35.8 S, 148°14.0 E), Port Hacking (PHB: 34°05.0 S 151°15.0 E), and North Stradbroke Island (NSI: 27°20.5 S 153°33.75 E) (Figure 1). The MAI station is situated 7.4 km off Maria Island, on the Tasmania east coast (depth 90 m) and is seasonally impacted by the southerly extent of the East Australian Current (EAC), which is a strong western boundary current [44]. PHB is located at the southern extent of the EAC separation zone (Figure 1) and 5.5 km offshore (depth 100 m). NSI is located north of Brisbane (depth 50 m), and is strongly influenced by EAC waters that originate in the Coral Sea [44]. Sampling at each time-series station comprised collection of bulk seawater samples for microbial analyses from mooring sites at near-monthly intervals (median days between sampling events; MAI: 34; PHB: 33; NSI: 32), with physico-chemical and Chl-a data (collectively termed environmental from here on) collected simultaneously for approximately 5.5 years (2012 – 2017), totaling 157 samples (MAI: 58; PHB: 47; NSI: 52; SI Table 1). Environmental variables measured at each site included temperature (°C), day length (hours), salinity (PSU), turbidity (NTU), Secchi disk depth (m), thermocline depth (m), dissolved silicate (umol/L), NOx (umol/L), phosphate (umol/L), ammonium (umol/L), and Chl-a concentration (mg/m^3^). Data were collected and analysed by IMOS [45, 46] (SI Table 1). Mixed layer depth (MLD) was estimated from temperature depth profiles (Australian National Mooring Network temperature and salinity data product at http://aodn.com) based on Condie and Dunn (2006) [47], and defined as the depth at which temperature decreased by 0.4 °C from the surface temperature (0-2 m depth).

**Figure 1:**
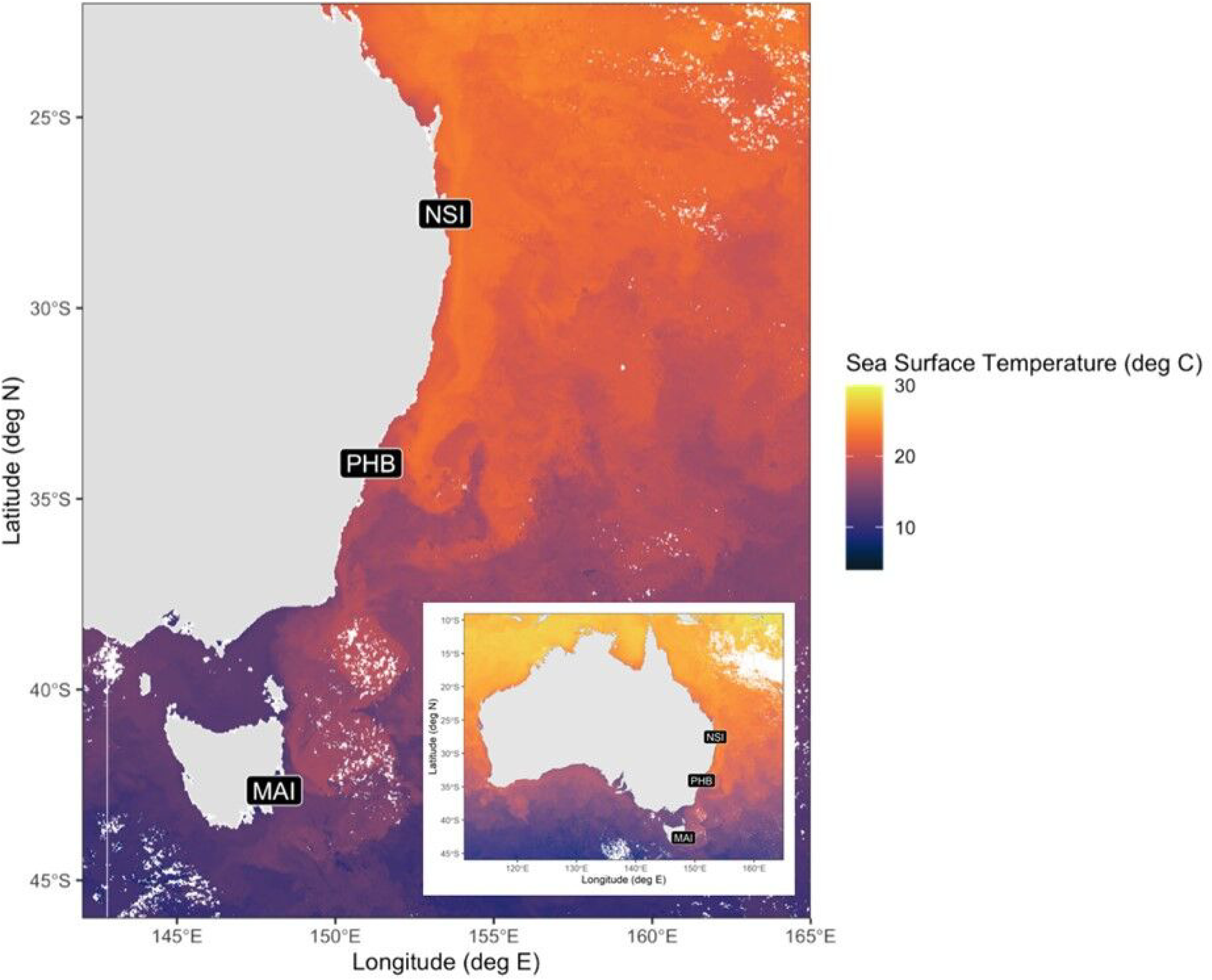
Map of sampling location. Inset figure shows the relative location of each reference station to the Australian continent.

### Sample collection, DNA extraction and amplicon sequencing

#### Sample collection and DNA extraction

Two liters of surface seawater were collected using Niskin bottles and transported on ice back to the lab. Samples were filtered through a 0.22 um pore Sterivex GP filter (Millipore, Massachusetts. Cat. # SVGPL10RC), which were then stored at −80 °C until processing. Filters were sent (on dry ice or in liquid nitrogen dewars) to the Commonwealth Scientific and Industrial Research Organisation Oceans & Atmosphere (CSIRO O&A) laboratories in Hobart, Tasmania for DNA extractions. Microbial DNA was extracted using standardized procedures as part of the Marine Microbes Program (https://data.bioplatforms.com/organization/pages/bpa-marine-microbes/methods) using a modified PowerWater Sterivex DNA Isolation Kit (MOBIO Laboratories) protocol. DNA isolation included incubating Sterivex filters for 1 hour on a horizontal vortex with 1.875 ml lysis buffer followed by a phenol:chloroform extraction.

#### Amplicon sequencing

The V1 – V3 regions of the 16S rRNA gene were PCR amplified using the bacteria-specific primers 27F (5’-AGRGTTTGATCMTGGCTCAG-3’) and 519R (5’-GWATTACCGCGGCKGCTG-3’) [49] with the following cycling conditions: 1-step using KAPA HiFi HotStart ReadyMix (Roche) comprised steps including 95 °C initial denaturation (3 min), with 35 cycles of 95 °C (30 s), 5 °C (10 s) and 72 °C 45 s), and a final elongation step at 72 °C (5 min). Amplicons were then purified using Ampure XP beads (Agencourt Bioscience Corporation) and sequenced on the Illumina MiSeq platform (Illumina, Inc., San Diego, USA) at the Ramaciotti Center for Genomics (UNSW, Sydney, Australia), with 300 bp paired reads.

### Bioinformatic processing

Raw fastq files were downloaded from BioPlatforms Australia (https://data.bioplatforms.com). Amplicon quality control and analysis was performed using DaDa2 [50]. In brief, primers were truncated using cutadapt [51] and reads were trimmed, denoised, merged, and chimeras removed using function removeBimeraDenovo (minFoldParentOverAbundance = 4) (full code provided https://github.com/martinostrowski/marinemicrobes/tree/master/dada2). Taxonomic classification of bacterial 16s rRNA ASVs was performed using a naïve Bayes classifier based on SILVA 138.1 and a bootstrap cut-off >50 % [52]. All ASVs which had a DaDa2 bootstrapped value < 50 at the taxonomic level were assigned to Kingdom unclassified.

The final bacterioplankton dataset analyzed in this study resulted from removing all sequences assigned to Kingdom unclassified, Archaea, Eukaryota, Chloroplast, and Mitochondria. The final step included filtering out low abundant ASVs with a total abundance across the entire dataset of less than 0.005 %. The plastid dataset containing all the Chloroplast sequences was used to assess the potential importance of phytoplankton on bacteria community assembly, and taxonomic assignment was made in a similar way as bacterioplankton ASVs with bootstrapped value < 50 % were trimmed and taxonomic identity called with naïve Bayes classifier using PhytoRef database [53].

### Statistical analysis

Datasets used in analysis included 1) environmental variables, 2) bacterial amplicon relative abundance, and 3) plastid amplicon relative abundance to represent the eukaryotic phytoplankton. In cases of missing environmental observations, values were imputed with rfImpute () from the randomForest package (version 4.6.14) [54]. Imputation was performed for each time-series independently. Environmental variables were mean centered unit variance standardized to reduce outlier influence. All analyses were performed using R version 3.6.1.

#### Temporal variability in environmental conditions

Temporal variability in environmental conditions was estimated by determining the mean dissimilarity within each time-series [37] based on the 11 environmental variables described above. Dissimilarity among samples based on environmental variables was calculated on Euclidian distance. Heterogeneity (*Ed*) was derived for each pair of samples within a time-series as follows:

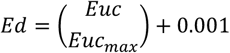

where *Euc* is the Euclidian distance between two samples within a time-series, *Euc*_*max*_ is the maximum distance observed across the entire dataset, and 0.001 is added to account for zero similarity among two samples. Mean *Ed* was then calculated within each time-series and heterogeneity compared using a Kruskal-Walls χ^2^ and dunn-post hoc pairwise tests across the three time-series.

#### Calculation of the biological interaction indices

Building on a framework introduced by Musters et al., (2019), we developed a metric to quantify the relative contribution of potential interactions among bacteria and phytoplankton in structuring patterns of bacterioplankton diversity. Our approach regresses biological predictors (bacterial and phytoplankton ASVs) against individual bacterial ASVs (response ASV). In the case of the bacterial interaction metric, the response ASV is removed from the predictor ASV dataset. There is no reason to expect species interactions will be linear or that ASV patterns are the result of a single predictor, therefore we extend the co-occurrence definition beyond simple pairwise co-occurrence to include more than a single bacterial ASV (or phytoplankton ASV) using a machine learning approach. In this way, we can identify non-linear abundance relationships of an individual ASV which may be due to the abundance of multiple organisms. We additionally identified the relationship of each bacterial ASV to a combination of environmental data (e.g, temperature). Therefore, we generated three datasets including bacteria-environment, phytoplankton-environment, and environment only in a series of steps (SI Figure 1) described below. The gradient forest [55] method was used to regress the large number of predictors to individual bacterial ASVs. Gradient forest is a modification of regression forest, which calculates an explained variation 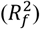 for each response ASV (SI Figure 1a,b). The approach then uses an out-of-bag prediction (OOB) similar to regression forest, but differs by defining the explained variation of each response and is calculated as follows:

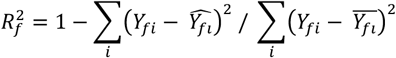

*Y*_*fi*_ is the abundance of the *ith* occurrence of ASV *f*, 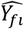 is the OOB prediction for the abundance of ASV *f* at the *ith* position, and 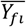 is the mean abundance of ASV *f*. Calculations were conducted using the gradientForest package (version 0.1.17) in R [56]. For each response variable we removed all predictor variables that did not significantly explain the abundance of the response ASV (SI Figure 1c). Significance of predictor variables in explaining proportional distribution of the response was calculated with rfpermute (version 2.1.81). Regression forest produces partial R^2^ values for each predictor of a response ASV. All partial R^2^ values for a response variable are summed to produce the total R^2^ of that response variable; therefore, we could sum the remaining significant predictor partial R^2^ values to determine the response R^2^ value (SI Figure 1d).

Next, we removed all bacterial response ASVs with an R^2^ value less than 0.3 (thus retaining >= 0.3) and summed the relative abundance of each retained ASV for each sample (i.e., sample 1 from NSI, sample 2 from MAI) (SI Figure 1e). Finally, the bacteria-bacteria metric was calculated as the abundance difference between the bacteria-environment and environment only relative abundance for each sample. The bacteria-phytoplankton metric was calculated as the difference between the phytoplankton-environment and environment only relative abundance (SI Figure 1f). The resulting value is the bacterial or phytoplankton interaction metric (SI Figure 1g). Our approach is similar to that described by Muster et al., (2019) with the addition of the significance calculation for partial R^2^ values and the identification of the relative abundance of ASVs from the dataset which are described by other bacterial or phytoplankton ASVs. Before the calculation of the metric, we performed a Hellinger transformation on predictors to reduce potential bias from highly abundant predictor ASVs. Also, to reduce computational time, we limited our analysis to include only ASVs which occurred in > 25 % of samples within a time-series.

#### Recurrent diversity patterns and environmental drivers

Alpha diversity was calculated as the effective number of ASVs [57] per sample, which is a measure of the number of equally common ASVs among samples and calculated as *e* ^*H’*^, where H’ is Shannon entropy. Beta-diversity was calculated using abundance-based Bray-Curtis dissimilarity for pairs of samples within time-series. Abundance based beta-diversity was calculated on Hellinger transformed data (square root of standardized species abundances). Bray-curtis similarity was used throughout and calculated as 1-dissimilarity score. To test for differences in intra-seasonal and inter-seasonal beta-diversity, samples were classified as intra-seasonal if sample pairs were from the same astronomical season; otherwise, samples were classified as inter-seasonal. A Kruskal-Wallis χ^2^ test was performed to test the null hypothesis of no difference among the means of time-series groups.

We then modeled predictor variables that best explained observed diversity (alpha and beta) across the three time-series. For these analyses, we included the biotic interaction metrics, as described above, to account for possible interspecific interactions and/or unobserved environmental factors. Multiple regression was run on alpha diversity using lm () and step () with the direction set to ‘both’, from the ‘stats’ package (version 3.6.1). Model selection was performed using AIC decrease. Distance-based modeling was performed to identify explanatory variables that best explained the distribution of beta-diversity [58], using a stepwise procedure. Model selection was based on the increase in adjusted R-squared with procedure ordiR2step () from the vegan package (version 2.5.6) [59] with direction = ‘both’ and permutations = 1000. We then performed variance partitioning [60], to identify the relative importance of the variables selected by the distance-based step procedure using the function varpart().

## Results and Discussion

### Environmental characteristics of the three oceanographic time-series sites

Environmental heterogeneity exhibited a latitudinally-defined gradient (Kruskal-Wallis χ^2^ _df = 2_ = 701.4, p < 0.01; Figure 2a, b, SI Figure 2a, b, c) across the three time-series stations, whereby Maria Island (MAI) had greater environmental heterogeneity than Pt Hacking (PHB) (Figure 2a; Dunn-test: p < 0.01) and PHB greater than North Stradbroke Is (NSI) (Figure 2a; Dunn-test: P < 0.01). At MAI (Figure 1; Lat 42°35.8 S; Lon 148°14.0 E) autumn and winter were characterized by a greater mixed layer depth (MLD; mean ± sd; 63.3 m ± 20.6), higher inorganic nutrient concentrations (SI Table 1; SI Figure 2b; NOx: 1.52 umol/L ± 1.42; phosphate: 0.216 umol/L ± 0/109) and lower temperature (SI Figure 2a; 15.1 °C ± 2.14), while spring and summer samples had higher Chl-a concentrations (SI Figure 2c; 0.572 mg/m^3^ ± 0.333). At PHB (Figure 1; Lat 34°05.0 S; Lon°151 15.0 E) spring and summer temperatures (SI Table 1; 20.2 °C ± 2.07) more closely track that of NSI than MAI (SI Figure 2a), while MLD (32.0 m ± 17.0), inorganic nutrient concentrations (SI Figure 2b; NOx: 1.00 umol/L ± 1.33; phosphate: 0.174 umol/L ± 0.10; silicate: 0.864 umol/L ± 0.749), and Chl-a concentration (SI Figure 2c; 0.67 mg/m3 ± 0.30) during winter were more similar to MAI. NSI (Figure 1; Lat 27°20.5 S; Lon 153°33.75 E) was distinguished by relatively high-water temperatures (SI Figure 2a; SI Table 1; 23.6°C ± 2.03), and relatively low concentrations of inorganic nutrients including phosphate (SI Table 1; 0.09 umol/L ± 0.04) and NOx (SI Figure 2a; 0.07 umol/L ± 0.13) concentrations. Winter samples at NSI were characterized by relatively high Secchi disk depth (20.0 m ± 5.33) while samples from the other three seasons were most distinguished by temperature. Thus, the three time-series exhibited distinct environmental conditions that range from MAI having the greatest environmental heterogeneity compared to other stations, PHB with relatively intermediate nutrient concentrations and high physical environmental heterogeneity and NSI having the least environmental heterogeneity. (324)

**Figure 2:**
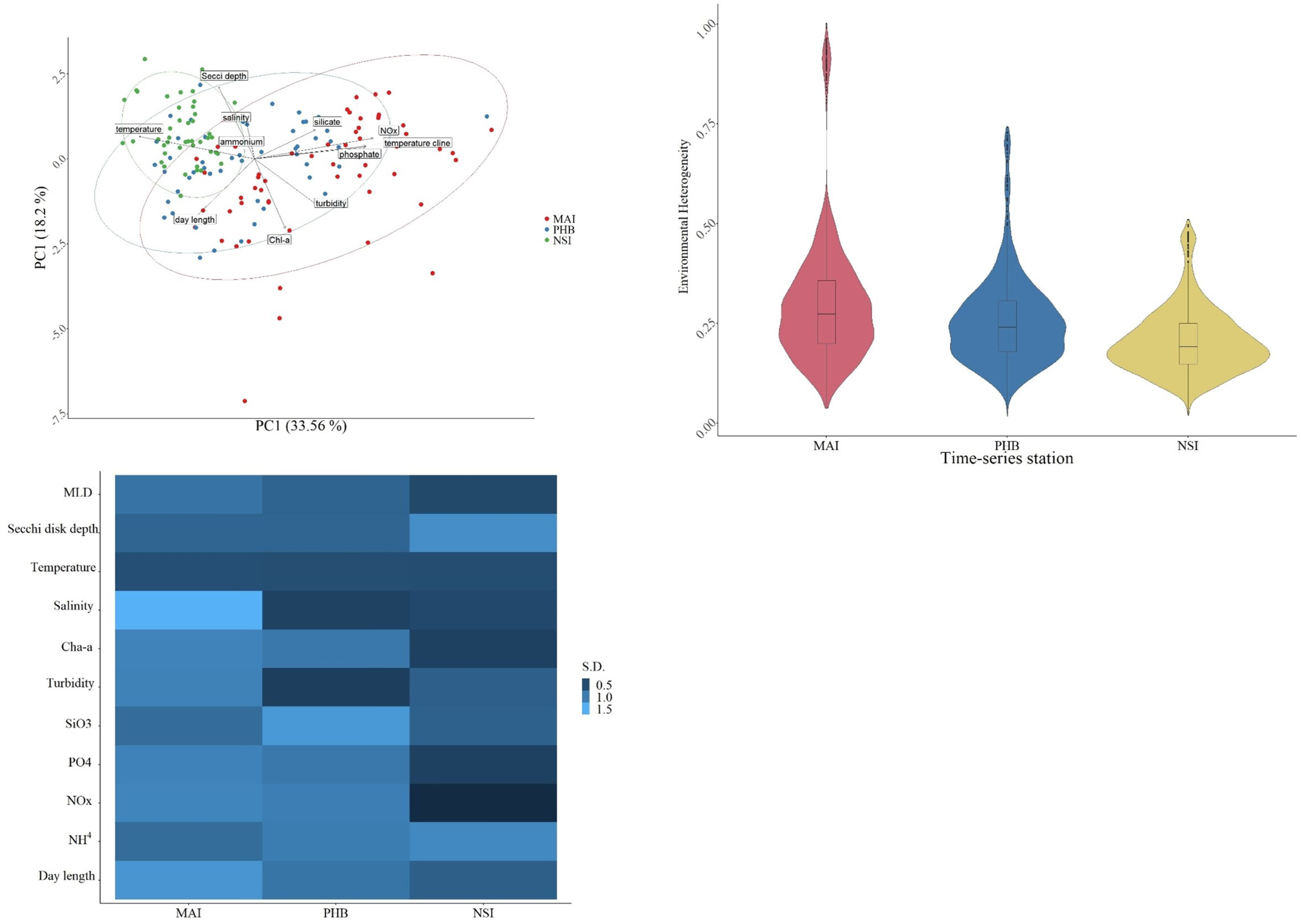
Environmental variability of each time-series. a) PCA biplot of first two dimensions discriminating samples to demonstrate environmental heterogeneity and is based on measured environmental variables. b) Distribution of environmental heterogeneity across time-series. c) Heatmap visualizing the relative heterogeneity, measured as standard deviation of environmental variables across time-series.

### Contrasting drivers of bacterioplankton alpha diversity patterns across time-series

The bacterioplankton datasets from the three reference stations had a varying number of observed ASVs (richness). Maria Island had the greatest number of total ASVs with 7608 (mean per sample 490.8 ± se 26.0), then Port Hacking with 7020 (414.1 ± 20.7) and North Stradbroke Island with 4843 (431.8 ±19.7). Richness of dataset ASVs corresponded with the total diversity of ASVs at each site where MI had the greatest alpha diversity (mean 122.14 ± se 7. 64), followed by PHB (116.68 ± 6.36) then NSI with the least (107.27 ± 4.07). The distribution of alpha diversity was, however, not significantly different among time-series (SI Figure 3; Kruskal-Wallis χ^2^ = 2.10, df = 2, p = NS). Temporal patterns in bacterioplankton diversity across the time-series sites provided evidence however, for varying degrees of seasonality among locations. Consistent yearly diversity patterns were observed at MAI, and this was less apparent or absent at PHB and NSI (Figure 3a). At MAI, bacterioplankton alpha diversity consistently peaked in the winter months and was lowest during spring, while at PHB, diversity peaked inconsistently across years. For instance, in 2012, the highest observed diversity at PHB was in winter, whereas in 2013, diversity peaked in autumn. At NSI, diversity peaks were not consistent across years. Collectively, these patterns infer that at MAI the principal factors regulating bacterioplankton alpha diversity are repeatable at seasonal scales, while at PHB and NSI, factors regulating bacterioplankton alpha diversity lack seasonal influences.

**Figure 3:**
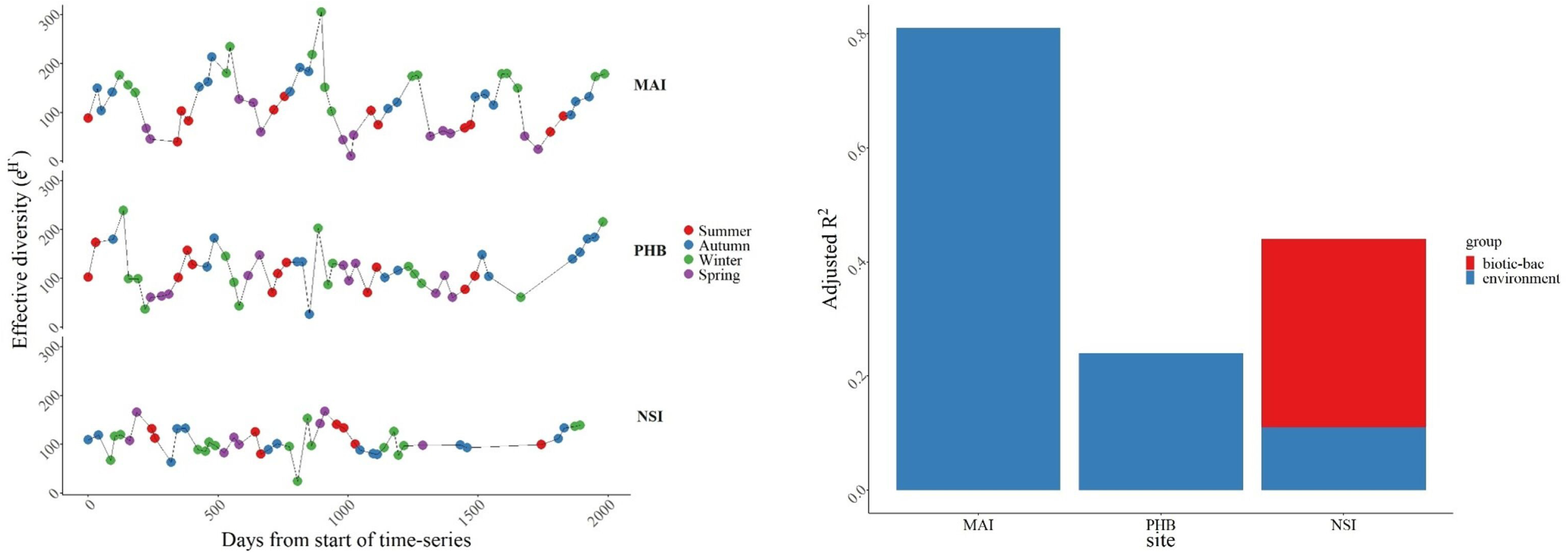
Patterns and drivers of alpha diversity across time-series. a) Scatterplot of alpha diversity through time for each time-series. X-axis is time from the start of the time-series and y-axis is effective diversity. Colors represent seasonal classification based on astronomical calendar. b) Contribution of environmental variables and the biological metric to explaining variation of alpha diversity through time at each time-series. The x axis displays the three time-series, and the y axis is the adjusted R2 score. Blue is the adjusted R2 from a multiple regression model of environmental variables only while red is the improved R2 when the biological metric is included in the model.

Environmental heterogeneity has been shown in other systems to be an important driver of bacterial diversity patterns [7, 61], therefore, given the different levels of environmental heterogeneity observed between locations, we predicted that the influence of environmental factors would become less apparent with decreasing environmental heterogeneity. At MAI, diversity patterns were predominately predicted by environmental factors (Figure 3b; Table 2; F _df = 4,53_ = 57.06, p < 0.001, Adj. R^2^ = 0.80). Day length had a strong inverse relationship with alpha diversity (Relative Importance = 0.62). Similar results have been reported for bacterioplankton richness patterns in the English channel time-series [25] which was also sampled at near monthly intervals. Therefore, day length may generally be an important predictor of high latitude bacterioplankton diversity. In addition, Chl-a was weakly associated with bacterioplankton alpha diversity suggesting a potential trophic mediation by phytoplankton (RI = 0.09). Similar trends were observed in the Antarctic where bacterioplankton alpha diversity was inversely related to Chl-a [26].

**Table 1:** Summary of imputed environmental variables. N is the number of samples in each time-series or the whole entire dataset. Min and max are the minimum and maximum observed values in the dataset. Mixed layer depth is estimated thermocline based on Condie and Dunn (2006).

|  | **Maria Island** | **Port Hacking** | **North Stradbroke Island** |
| --- | --- | --- | --- |
|  | **(N=58)** | **(N=52)** | **(N=47)** |
| **Temperature (°C)** |  |  |  |
| Mean (SD) | 15.1 (2.14) | 20.2 (2.07) | 23.6 (2.03) |
| Median [Min, Max] | 14.5 [11.9, 20.3] | 20.1 [16.8, 24.5] | 23.4 [20.4, 27.6] |
| Missing | 9 (15.5 %) | 3 (5.8 %) | 6 (12.8%) |
| **Day length (hr)** |  |  |  |
| Mean (SD) | 11.7 (2.06) | 11.9 (1.53) | 11.8 (1.14) |
| Median [Min, Max] | 11.5 [9.04, 15.3] | 11.8 [9.89, 14.4] | 11.5 [10.4, 13.9] |
| Missing | 0 (0%) | 0 (0%) | 0 (0%) |
| **Salinity (PSU)** |  |  |  |
| Mean (SD) | 35.2 (0.801) | 35.5 (0.193) | 35.5 (0.234) |
| Median [Min, Max] | 35.3 [29.3, 35.7] | 35.5 [34.7, 35.7] | 35.5 [34.3, 35.8] |
| Missing | 9 (15.5%) | 3 (5.8%) | 6 (12.8%) |
| **Turbidity (NTU)** |  |  |  |
| Mean (SD) | 0.388 (0.214) | 0.115 (0.0640) | 0.131 (0.139) |
| Median [Min, Max] | 0.285 [0.146, 1.10] | 0.102 [0.0582, 0.450] | 0.0878 [0.0108, 0.705] |
| Missing | 9 (15.5%) | 8(15.4%) | 8 (17.0%) |
| **Secchi disk depth (m)** |  |  |  |
| Mean (SD) | 16.5 (3.40) | 15.2 (3.44) | 20.0 (5.33) |
| Median [Min, Max] | 16.5 [9.00, 24.0] | 16.0 [9.00, 24.0] | 19.0 [9.00, 34.0] |
| Missing | 1(1.7%) | 3(5.8%) | 0(0%) |
| **Silicate (umol/L)** |  |  |  |
| Mean (SD) | 0.660 (0.487) | 0.865 (0.749) | 0.586 (0.414) |
| Median [Min, Max] | 0.600 [0, 2.00] | 0.800 [0, 3.90] | 0.500 [0, 2.10] |
| Missing | 7(12.1%) | 7(13.5%) | 5(10.6%) |
| **NOx (umol/L)** |  |  |  |
| Mean (SD) | 1.52 (1.42) | 1.00 (1.33) | 0.0679 (0.125) |
| Median [Min, Max] | 1.85 [0, 5.20] | 0.500 [0, 7.00] | 0 [0, 0.500] |
| Missing | 7(12.1%) | 8(15.4%) | 5(10.6%) |
| **Phosphate (umol/L)** |  |  |  |
| Mean (SD) | 0.216 (0.109) | 0.174 (0.0975) | 0.0934 (0.0365) |
| Median [Min, Max] | 0.210 [0.0200, 0.480] | 0.158 [0.0300, 0.650] | 0.0903 [0, 0.190] |
| Missing | 7(12.1%) | 7(13.5%) | 5(10.6%) |
| **Ammonium (umol/L)** |  |  |  |
| Mean (SD) | 0.164 (0.336) | 0.350 (0.412) | 0.293 (0.468) |
| Median [Min, Max] | 0.0772 [0, 2.40] | 0.231 [0.0300, 2.24] | 0.125 [0, 2.60] |
| Missing | 7(12.1%) | 10(19.2%) | 6(12.8%) |
| **Chl-a (mg/m<sup>3</sup>)** |  |  |  |
| Mean (SD) | 0.572 (0.333) | 0.669 (0.298) | 0.300 (0.111) |
| Median [Min, Max] | 0.526 [0, 1.62] | 0.617 [0.201, 1.41] | 0.293 [0.0810, 0.637] |
| Missing | 6(10.3%) | 14(26.9%) | 9(19.1%) |
| **Mixed-layer depth (m)** |  |  |  |
| Mean (SD) | 63.3 (20.6) | 32.0 (17.0) | 31.4 (9.92) |
| Median [Min, Max] | 74.0 [21.0, 86.0] | 27.3 [11.0, 81.0] | 31.0 [13.0, 57.0] |
| Missing | 8(13.8%) | 4(7.7%) | 3(6.4%) |

**Table 2:**
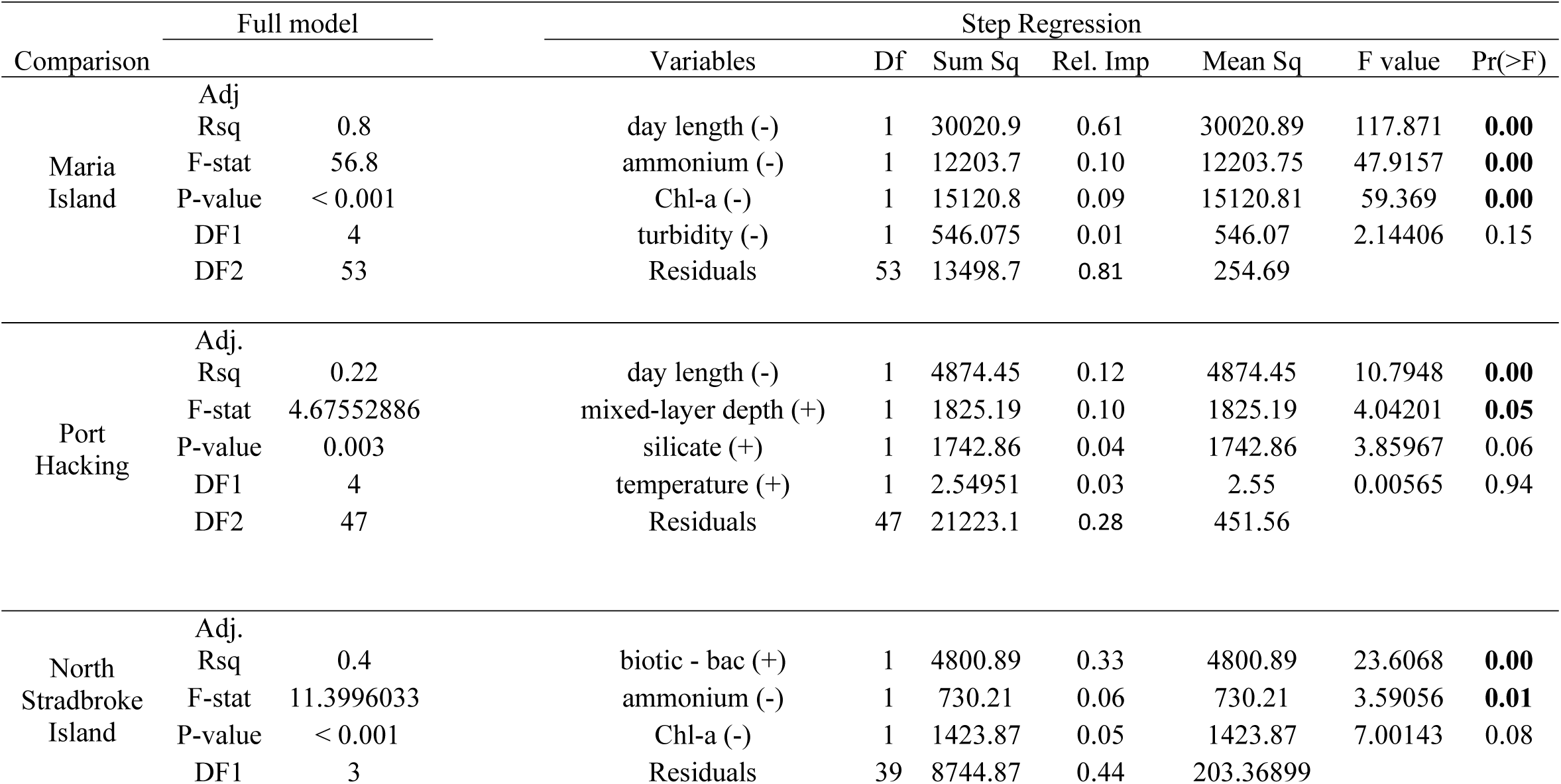
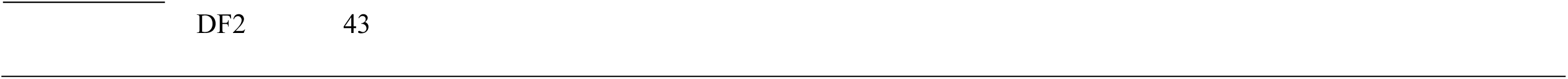
Linear model results for predictor variables that showed the best relationship with patterns of bacteria alpha diversity through time. The full model is the results of all variables. Individual variables are the result of step regression. The (+) and (-) indicate the direction of relationship between variables and alpha diversity. The biological metric is the interspecific interaction metric. The explained variation (Exp.var) is the result of partitioning the variable sums of squares. DF = degrees of freedom; Sum Sq = Sums of square; Rel. Imp = Relative importance based on CAR variance partitioning; Mean Sq = Means of the square. Significance represented by bold font if p < 0.05.

At PHB, where there were lower levels of seasonal heterogeneity in environmental conditions (Figure 2a), bacterioplankton diversity was influenced by mixed layer depth, but total explained variation for alpha diversity was quite low (Figure 3b; Table 2; F df = 5, 46 = 4.29, p = 0.003, Adj. R^2^ = 0.24). The dominate environmental factors included day length (RI = 0.12) which was inversely correlated with alpha diversity patterns while MLD depth (RI = 0.10) was positively correlated. The high amount of unexplained variation may suggest other unmeasured environmental factors (e.g., dissolved organic carbon) more strongly influence alpha diversity. Alternatively, EAC driven dispersal processes, which have been shown to influence bacterioplankton occurrences at PHB [48], may also be a dominant contributor to alpha diversity at the monthly time-scale interval. Dispersal is a fundamental ecological process [62] and can become important in structuring bacterioplankton diversity when environmental heterogeneity is low or when dispersal rates are high enough to over-shadow the effects of other ecological processes [7].

For NSI temporal bacterioplankton alpha diversity was not consistent with astronomical seasons, but total variation could be explained to a relatively high level (Figure 3b; Table 2; F df = 3,43 = 11.18, p < 0.001, Adj. R2 = 0.40). Interestingly, and in contrast to the other two locations, biotic interactions were the main predictors of alpha diversity at this location. Bacteria-bacteria interactions specifically, were positively correlated with alpha diversity patterns and contributed a large portion of the total predicted variation (Figure 3b; partial R^2^ = 0.33). Ammonium was also important in predicting alpha diversity and was inversely correlated (RI = 0.06) with diversity patterns. Therefore, potential interspecific interactions may be important drivers of alpha diversity patterns at this sub-tropical time-series [33].

Across the three time-series the amount of variance that could be explained by environmental factors corresponded with trends in environmental heterogeneity. The largest contribution of environmental variables to explaining alpha diversity distribution was at MAI (80 %), while an intermediate amount could be explained at PHB (22 %) and the least at NSI (10 %) (Figure 3b). However, the total explained variation did not correspond with trends in environmental heterogeneity. The lowest latitude site NSI which had the lowest environmental heterogeneity, had the second largest total explained variance, driven by a large contribution of biotic predictors (24 %). This location had the warmest temperatures and the lowest inorganic nutrient concentrations of our study locations (Figure 2b), and under these conditions trophic mediation, such as facilitation by Prochlorococcus and Synechococcus groups can drive bacterioplankton succession through primary productivity [63]. Biotic interactions at MAI or PHB were not important predictors of alpha diversity across the temporal scale analyzed here (median 34 days), however is likely an important contributor when higher resolution time-series are considered [64]. For instance, Luria et al (2016) monitored bacterioplankton diversity in Antarctic waters across 1-2-week intervals and found richness was driven phytoplankton blooms; therefore, potentially demonstrating importance of scales in distinguishing among dominate ecological drivers of diversity patterns.

### Contrasting drivers of recurrent beta diversity patterns across time-series

Bacterioplankton beta diversity (ratio of regional: local diversity) at each of the three reference stations exhibited seasonal trends, where intra-seasonal samples (samples from the same season) had greater observed similarity (ie. lower beta diversity; 0: dissimilar; 1: highly similar) than inter-seasonal samples (Figure 4a, Supplemental Figure 4). At MAI, the mean intra-seasonal Bray-Curtis (BC) score of 0.50 (± 0.005 SE) was significantly greater than the inter-seasonal score (0.59 ± 0.13; t-test _df = 749.45_ = 16.00; p < 0.001). Similarly, at PHB the intra-seasonal similarity (0.56 ± 0.006) was significantly greater than the inter-seasonal similarity (BC = 0.63 ± 0.003; Supplemental Figure 4; t-test _df = 528.48_ = 10.32; p < 0001). NSI had the lowest intra-seasonal mean BC among the time-series at 0.47 ± 0.005 which was also significantly different than the inter-seasonal mean BC of 0.51 ± 0.003 (Supplemental Figure 4a; t-test _df = 473.94_ = 6.00; p < 0.001). Therefore, at all locations bacterioplankton communities from a given season were more similar to those from the same season in different years, than to those that were closer in time, but different in season. These results suggest ecological processes that structure bacterioplankton communities are recurrent at a given time of across years, and this occurs across despite.

**Figure 4:**
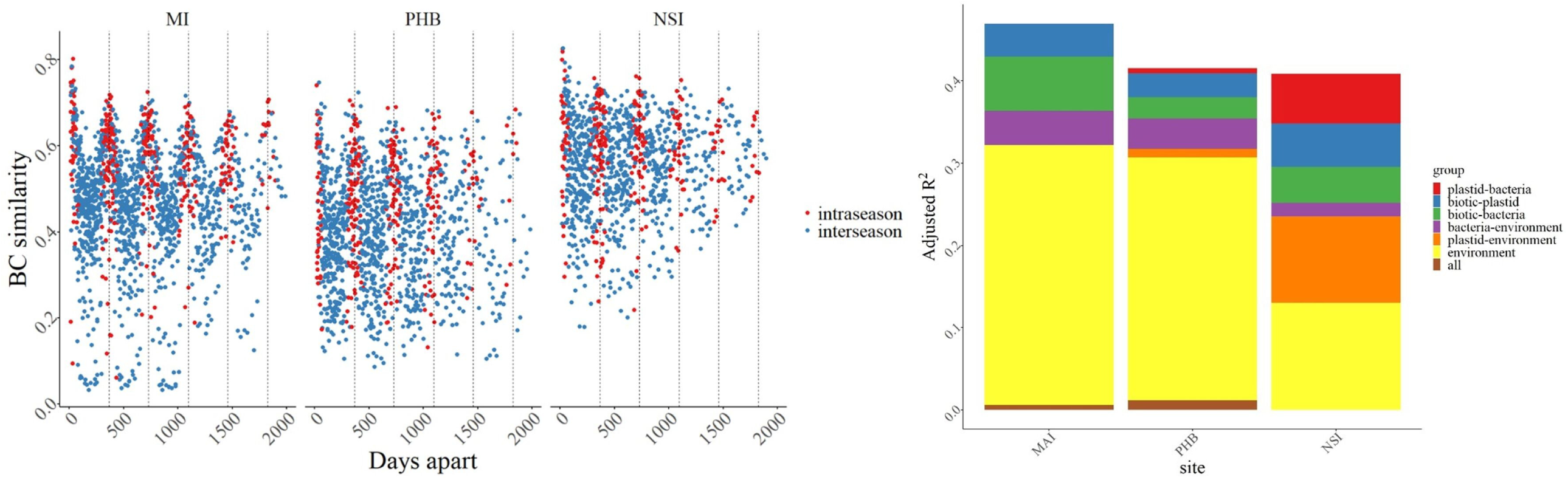
Beta diversity patterns and contribution of deterministic drivers. a) Time (in days) between sampling points along X-axis and Bray-Curtis dissimilarity (BC) scores along the Y-axis. BC = 0, entirely the same; BC = 1 entirely different. Colors represent the category of sample; blue = inter-seasonal (two samples from different seasons), red = intra-seasonal (two samples from the same season). Dotted vertical line breaks spaced at 365 days to show length of time series. b) Contribution of environmental and the biological metric to explaining variation of beta diversity through time across each time-series. Colors correspond to the amount of variation attributed to several ecological processes derived from variance partitioning procedure.

Like alpha diversity, beta diversity is also expected to increase with increasing environmental heterogeneity [65, 66] under the assumption that greater variability in environmental factors will result in an increased number of niches for organisms to occupy [67]. We therefore predicted beta diversity would be greatest at MAI and lowest at NSI. This pattern, however, was not observed and rather the greatest mean beta diversity was observed at PHB (Supplemental Figure 4b; mean ± SD; 0.61 ± 0.11), followed by MAI (0.57 ± 0.12) and NSI (0.50 ± 0.09; Kruskal-Wallis χ^2^ _df = 2_ = 690.3, p < 0.05). These results suggest that environmental variability is not entirely responsible for bacterioplankton composition, suggesting other ecological processes, such as biotic processes are important for structuring beta diversity.

Therefore, we investigated the key variables driving beta diversity patterns and determined their relative contributions to these patterns. Our results indicate that different deterministic processes govern patterns of beta diversity across the three locations (Figure 4b). Variables that best modelled beta diversity at MAI included day length, temperature, bacterial abundance, phytoplankton abundance, turbidity, and Secchi disk depth (Adj. R^2^ = 0.20, 0.13, 0.06, 0.04, 0.02, < 0.01, respectively; Table 3; F= 11.35, df = 6, 51, p = 0.001). Variance partitioning showed environmental variables had the greatest effect (32 % of partitioned variation; Figure 4b) influencing beta diversity patterns at MAI followed by bacterial interactions (7 %), while phytoplankton contributed 4 %. Together, the biotic interactions explained approximately 11 % of the total partitioned variation. There was 4 % of variance contributed by bacteria-environment overlap, suggesting a potential role of environmentally mediated bacterial influence. These results match with alpha diversity patterns where environment was the key drivers, demonstrating the importance of environment fluctuation in structuring bacterioplankton diversity. Beta diversity however had some influence by biotic factors, while alpha diversity was only predicted by environmental factors, potentially suggesting that biotic processes may facilitate the presence or absence of particular bacterioplankton groups, rather than diversity at a particular time point.

**Table 3:**
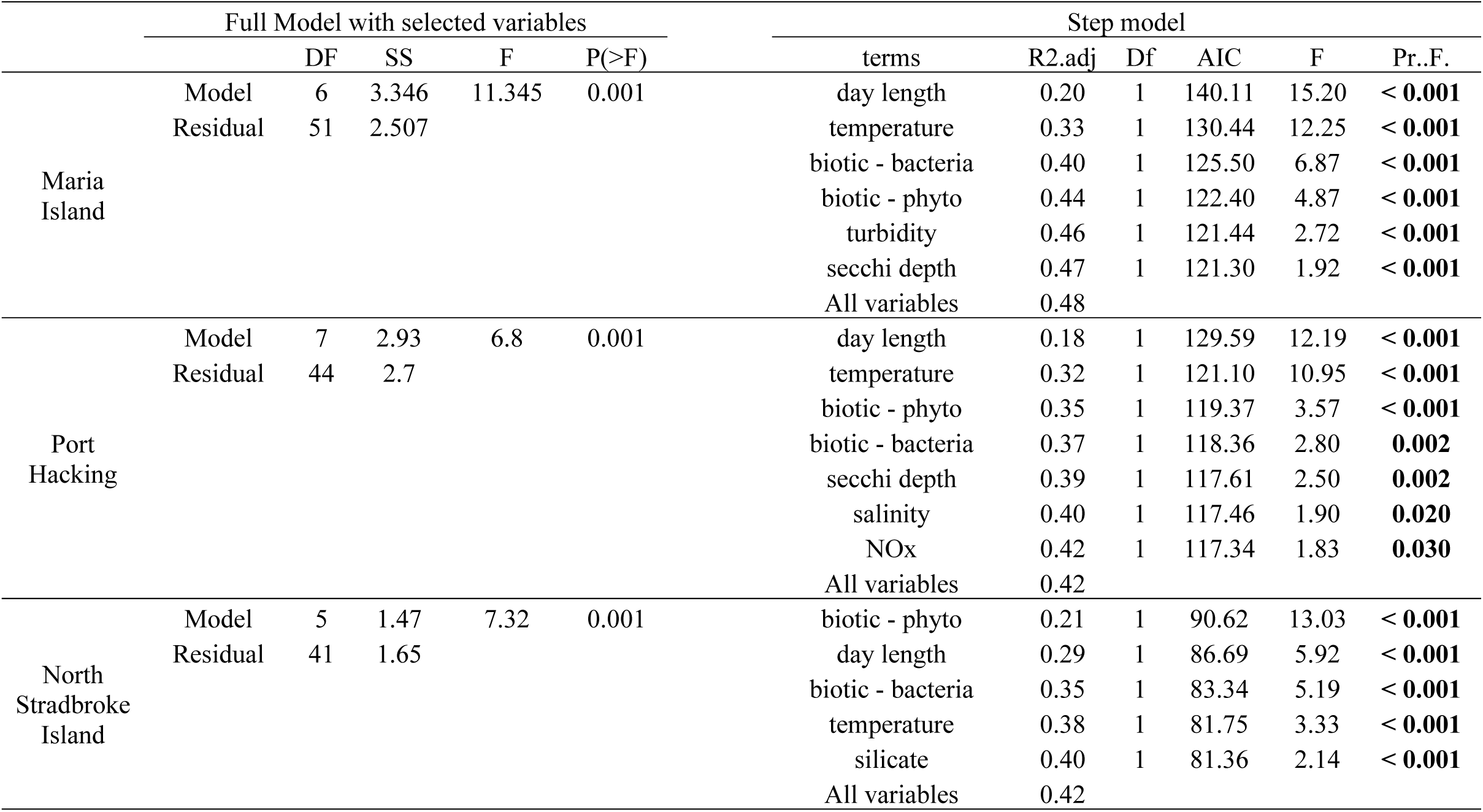
Distance-based linear model results for predictor variables that showed the strongest relationship to patterns of beta diversity through time. The biological metric is the interspecific interaction metric. Phyto is in reference to the phytoplankton biological metric. The full model informs on the global test for all selected variance and the step model shows the results for individual chosen variables. All variables are the model results when all variables are included. Df = degrees of freedom; SS = sums of squares; AIC = Aikaike information criteria.

At PHB, environmental factors also had the largest contribution to beta diversity. Important variables included day length and temperature (Adj R^2^ = 0.18, 0.14, respectively; Table 3; F= 6.8, p = 0.001). Collectively, the environmental factors accounted for 30 % of the total partitioned variation (Figure 4b) while biotic interaction (bacteria and phytoplankton) only accounted for 5 % of the total variation. Environmental overlap with bacteria (4%) and phytoplankton (1 %) accounted for 5 % of the variation. These results are similar to alpha diversity in that environment was a key contributor to observed patterns. Interestingly, environmental contribution was similar to the amount contributed at MAI, however total explained variance was lower due to the lower contribution of biotic influence at PHB.

A key finding in this study was that at NSI, biotic predictors played a much greater role in defining beta diversity relative to the other two locations. Phytoplankton abundance was found to be the most important factor contributing to bacterioplankton beta diversity variation (R^2^ = 0.21; Table 3; F= 7.32, df = 5, 41, p = 0.001). Biotic factors accounted for the largest amount of partitioned variation at 15 % (Figure 4b; phytoplankton-bacteria: 6 %; phytoplankton only: 5 %; bacteria only: 4 %) while environmental factors only accounted for 13 % of the variation. There was a large amount of variation accounted for due to overlapping components, including phytoplankton-environment (11 %) and bacteria-environment (2 %) (Figure 4b; green segment). Based on the high observed influence of phytoplankton abundance at NSI and high overlapping variance between phytoplankton and the environment, we posit that the environment may indirectly drive bacterioplankton beta diversity through influencing the phytoplankton. These results are similar to those observed for alpha diversity patterns, where biotic predictors were also the most important contributor. Interestingly, the main biotic contributor varied across the two diversity measures, where phytoplankton was the most importance for beta diversity while for alpha diversity, bacteria was the predominate drivers. Thus, trophic links are important to structuring bacterioplankton diversity in a dynamic manor at NSI.

Together these results show that patterns of beta diversity are not shaped by environment alone, but rather a combination of environment and potential biotic interactions and that the relative importance of these can vary across locations. Interestingly, the importance of biotic interactions negatively corresponded with beta-diversity, such that the total contribution by biotic factors was greatest at NSI where beta-diversity was lowest, while PHB had the highest beta diversity and was least influenced by biotic predictors. These results potentially signal a stabilizing effect on the community against environmental fluctuation that biotic interactions can promote [68]. Also, in contrast to predictions, the relative contribution of deterministic processes did not entirely correspond with changes with environmental heterogeneity, as the relative contribution of environmental factors were similar at MAI and PHB, however at NSI where the lowest level of environmental heterogeneity occurred, biotic processes were the predominate deterministic driver. Interestingly, biotic influence on bacterioplankton diversity was found at all locations, suggesting previously overlooked factors driving temporal succession of bacterioplankton.

## Concluding remarks

Here, we demonstrate that temporal patterns in marine bacterioplankton diversity are structured by different inherent deterministic processes according to location, which tracks latitudinal differences that may be the result of variation in environmental heterogeneity. The most ‘environmentally stable’ site, which was characterized by the least seasonality displayed patterns in bacterioplankton alpha and beta diversity which were in contrasted to the site with highest levels of seasonality in environmental conditions. Bacterioplankton diversity is the consequence of multiple interacting processes including filtering by environmental factors and biotic interactions [8, 69]. Partitioning the effects of environmental versus potential biotic influence is an important distinction as ecological theory predicts ecosystem function is linked to the processes that structure community diversity patterns [2, 70, 71]. Therefore, to accurately forecast ecosystem function, it is necessary to 1) distinguish among processes that give rise to bacterioplankton diversity and 2) identify how these processes change through space and time. This is heightened as climatic conditions are changing rapidly which can alter the balance between biotic and environmental deterministic processes [72]. However, until now no framework has been applied to bacterioplankton to identify the importance of potential biotic interactions relative to environmental factors driving total diversity patterns. Patterns of seasonality for both alpha and beta diversity observed in our study are consistent with diversity patterns from three well studied time-series, where the high latitude English Channel exhibited the highest degree of seasonality in diversity patterns, the mid-latitude SPOTS with intermediate diversity patterns and the low latitude HOTS with absent seasonal diversity patterns [73]. Therefore, processes driving diversity patterns along the latitudinal gradient may be general, and this study provides insight on potential drivers of this trend. Importantly, results shed insight on why some studies have identified environmental factors as having significant influence over bacterioplankton diversity [26], while others have concluded biotic processes play a stronger role in driving bacterioplankton diversity patterns [25, 69]. Predicting how biogeochemical processes will respond under future climate change scenarios requires insight to the microbial composition present, and therefore microbial diversity patterns.

## Acknowledgements

This project was supported by the New South Wales state government through the Research Attraction and Acceleration Program (). For samples collected prior to mid-2015, we would like to acknowledge the contribution of the Australian Marine Microbial Biodiversity Initiative (AMMBI) in the generation of the data used in this publication. AMMBI was funded by Australian Research Council awards DP0988002 to Brown & Fuhrman, DP120102764 to Seymour, Brown & Bodrossy, DP150102326 to Brown, Ostrowski, Fuhrman & Bodrossy, the Environmental Genomics Project from CSIRO Oceans and Atmosphere and a CSIRO OCE Science Leader Fellowship to Bodrossy. AMMBI was also supported by funding from the Integrated Marine Observing System (IMOS) through the Australian Government National Collaborative Research Infrastructure Strategy (NCRIS) in partnership with the Australian research community. For samples collected after mid-2015 we would like to acknowledge the contribution of the Marine Microbes consortium in the generation of data used in this publication. The Marine Microbes project was supported by funding from Bioplatforms Australia and the Integrated Marine Observing System (IMOS) through the Australian Government National Collaborative Research Infrastructure Strategy (NCRIS) in partnership with the Australian research community.

## Compliance with ethical standards

### Author contributions

MPD, JS, MAD, PS, and MO conceived and designed study; MPD, MO, and AB performed data processing; MPD performed statistical analysis; JS, MAD, MB, LB, JVDK assisted with sample collection; MPD wrote the manuscript with contributions from JS and MAD; and all authors edited the manuscript.

## Figure and Table Captions

**Supplemental Figure 1:** Biotic interaction index diagram. Each dataset is comprised of samples and corresponding ASVs as proportional abundance and standardized environmental variables (A). One ASV is separated (response variable) from other ASVs and environmental data (predictor variables) (B). Only ASV were included as response variables. A predictor by response matrix is returned with the partial R2 contributed to each response ASV (C) and the total R2 was calculated by summing partial R2 that were identified as significant (D). ASV’s with a total R2 less than 0.3 were removed from each sample and the relative proportion of each ASV was summed to get sample total (E). Random forest was run three times to obtain the sample total across a Bacteria-Environment dataset, Phytoplankton-Environment dataset and Environmental only dataset. Sample totals from biotic-environment (bacteria or phytoplankton) and sample totals from Environmental were subtracted (F) to obtain Biotic only dataset (Bacterial or Phytoplankton) indices (G).

**Supplemental Figure 2:** a) Temperature through time at each reference station. b) NOx concentration through time at each reference station. c) Average Chl-a concentration for each month for the three time-series.

**Supplemental Figure 3:** Alpha diversity plotted across each time-series. Boxplots represent the distribution of effective diversity scores at each time-series. Line indicates median score with either side representing the 2nd and 3rd quantile score distributions.

**Supplemental Figure 4:** a) The X-axis is the seasonal category, and the Y-axis is distribution of BC dissimilarity scores (0 = completely dissimilar, 1 = highly similar). Boxplot comparisons are partitioned into the three time-series. b) Boxplot of beta-diversity variation at each site.

